# Bacterial diversity and population dynamics during the fermentation of palm wine from Guerrero Mexico

**DOI:** 10.1101/480038

**Authors:** Fernando Astudillo-Melgar, Adrián Ochoa-Leyva, José Utrilla, Gerardo Huerta-Beristain

**Author notes:** Co-corresponding authors: José Utrilla, Gerardo Huerta-Beristain.

## Abstract

Palm wine is obtained by fermentation of palm tree sap. In the Pacific coast of Mexico, palm wine is called Tuba and it is consumed as a traditional fermented beverage. Tuba has empirical applications such as an auxiliary in gastrointestinal diseases and a good source of nutrients. In the present study, a next-generation sequencing of the V3-V4 regions of the 16S rRNA gene was employed to analyze bacterial diversity and population dynamics during the fermentation process of Tuba, both in laboratory controlled conditions and in commercial samples from local vendors. Taxonomic identification showed that *Fructobacillus* was the main genus in all the samples, following by *Leuconostoc*, *Gluconacetobacter*, *Sphingomonas,* and *Vibrio*. Alpha diversity analysis demonstrated variability between all the samples. Beta diversity clustered the bacterial population according to the collection origin of the sample. Metabolic functional profile inference showed that the members of the bacterial communities may present the vitamin, antibiotic and antioxidant biosynthesis genes. Additionally, we further investigated the correlation between the predominant genera and some composition parameters of this beverage. This study provides the basis of the bacterial community composition and functionality of the fermented beverage.

## Introduction

A wide variety of fermented food products such as yogurt, alcoholic beverages, bread, and sauces are produced worldwide. During the production process of these fermented foods, different microorganisms contribute to the organoleptic and biochemical characteristics (Tang et al., 2017). Recent studies in fermented food have shown that microbial ecology aspects such as diversity, their spatial distribution, and ecological interaction, have a strong influence on metabolic production and chemical composition (Escalante et al., 2015). Bacterial consortia interactions in fermented foods promote the process of polymer degradation and production of compounds of interest such as alcohol, aromatics, organic acids such as acetate, lactate among other metabolites that contribute to functional and organoleptic properties (Tamang et al., 2016).

Palm wine is a traditional beverage made using the sap collected from palm trees. It is consumed in different parts of the world, in Africa it is known as “legmi”, in South India as “kallu”, while in Borneo it has the names of “bahar” and “goribon” (Velázquez-Monreal et al., 2011). The differences among these beverages are the production process, the coconut tree species and the plant part where the sap is collected (Santiago-Urbina and Ruíz-Terán, 2014). In Mexico, several traditional fermented beverages are produced such as pulque (Escalante et al., 2016), pozol (Díaz-Ruíz et al., 2003), and Tuba (De la Fuente-Salcido et al., 2015). Tuba was brought to Mexico by Philippine influence during the Spanish colonial period. This beverage is produced in the southern Pacific coast of Mexico (Guerrero, Colima, Michoacan states). It is obtained from the sap of the inflorescences of *Cocos nucifera* L and it is consumed as a traditional beverage. Tuba is empirically used by locals as an aid in gastrointestinal problems and as a rehydration drink (De la Fuente-Salcido et al., 2015; Velázquez-Monreal et al., 2011).

The importance of bacteria in fermented foods has promoted the application of different strategies to analyze the bacterial diversity and role during their elaboration process. The use of massive sequencing technologies together with recent bioinformatics methods, such as QIIME for diversity analysis (Caporaso et al., 2010; Navas-Molina et al., 2015) and PICRUSt for functional inference (Langille et al., 2013), have increased the taxonomic and functional information of uncultured bacterial communities in different ecosystems (Filippis et al., 2017). However, those methods have been used mainly in projects such as Human Microbiome and Earth Microbiome (Creer et al., 2016). Nevertheless, in the food area, the applications of them are limited. Some studies in traditional Asian liquors and sauces have established a correlation between microbial diversity and organoleptic properties, increasing the information about bacterial communities in Asian products such as Yucha (Tang et al., 2017; Zhang et al., 2016).

Here, we study the fermentation profile, population dynamics and bacterial diversity of Tuba produced in the Guerrero coast of Mexico. We sampled Tuba production fermented under controlled conditions and compared it with local commercial Tuba. Using 16S amplicon sequencing and metabolic characteristics, we were able to analyze the diversity and infer functionality of bacterial communities present during Tuba production. This work provides a basis for the further functional characterization of Tuba in its production process, probiotic potential and other functions as antibiotic and antioxidant biosynthesis.

## Material and Methods

### Sap and commercial sample collection

Sap samples to ferment were collected on 14/07/2016 from three visibly healthy palm trees of the same palm producer in a rural area in Acapulco, Guerrero, Mexico. Commercial Tuba samples were obtained on 16/08/2016 from four different artesian producers in Diamante zone from Acapulco, Guerrero, Mexico (**Figure 1**). The climatological conditions, on the samples collection site at the sampling day, were similar with a temperature of 30-32°C, 87-89% of humidity, a pressure of 0.996 atm and a weather of light rain. Samples were transported in sanitized coolers to the laboratory for fermentation and analysis. The sap from palm trees was tagged with the following code a “P” followed by the number of the palm tree and “T” which means the fermentation time (i.e. P1T0). Commercial samples were tagged using the letter L followed by a consecutive number that symbolizes the number of the establishment where each sample was obtained. The total samples analyzed were 16.

**Figure 1.**
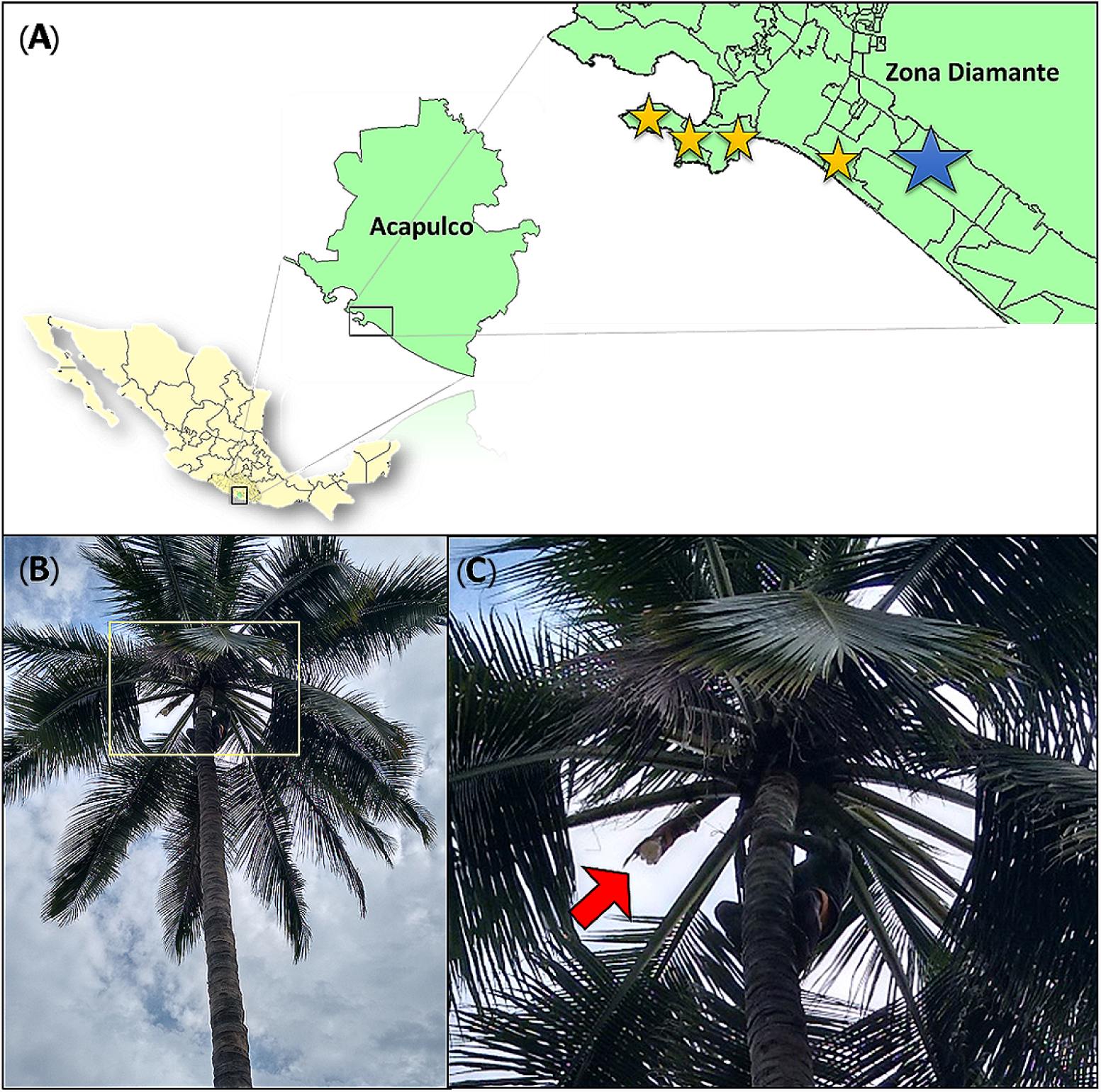
Sample collection. A) Sampling sites. The yellow stars represent the location of the four commercial establishments (commercial samples) and the blue star show the area where sap samples for the laboratory controlled fermentation were obtained. B) *Cocos nucifera* L (palm tree). Yellow square signaling sap collection zone. C) Sap collection zone. Red arrow indicate the palm structure (inflorescence) where the sap is collected.

### Fermentation in laboratory controlled conditions

Each sap sample from a different palm (100 mL as working volume) was fermented in four 250 mL Erlenmeyer flasks corresponding to 0, 12, 24 and 35 hours of fermentation. The fermentation was carried out as close to the commercial process as possible, we only used the native microbial community already present in the sap samples as a starter, no other inoculum was used. They were incubated at 30°C and 100 rpm of shaking speed in an orbital incubator. Samples were centrifuged (4000 rpm x 15 min) and the pellets were used for DNA extraction, while the supernatants were stored at −20°C for further analysis.

### Sucrose, glucose, fructose, xylose water-soluble proteins, acetic acid, ethanol, and pH

Sugars, organic acids, and ethanol from laboratory fermented and commercial samples were quantified using two HPLC methods following column manufacturer conditions. Glucose, fructose, sucrose, and xylose were quantified using an Aminex HPX-87P (Biorad) column with an IR detector. Acetate and ethanol concentrations were quantified using Aminex HPX-87H (Biorad) column and a UV 210 nm detector. Water-soluble proteins were measured by the Bradford method modified by Fernández & Galván, 2015. The pH was measured using a potentiometer with 1 mL of the sample.

### 16S amplicon library preparation and sequencing

The DNA extraction from all the samples was performed using the ZR Soil Microbe DNA MiniPrep™ kit (Zymo Research) according to the manufacturer protocol. The DNA was quantified using Qubit Fluorometric Quantitation (Thermo Fisher Scientific). 12.5 ng of total DNA was used for PCR of amplicons of the V3-V4 regions of the 16S rRNA ribosomal gene (F 5’-TCGTCGGCAGCGTCAGATGTGTATAAGAGACAGCCTACGGGNGGCWGCAG-3’ and R 5’-GTCTCGTGGGCTCGGAGATGTGTATAAGAGACAGGACTACHVGGGTATCTAATCC-3’) as described by the Illumina Protocol. All the PCR products were purified (AMPure XP beads - Illumina products) and quantified (Qubit). Finally, all the libraries were sequenced by Illumina MiSeq.

### Bioinformatics and Statistical analysis

The sequences were analyzed using QIIME (Quantitative Insights Into Microbial Ecology) version −1.9.1software (Caporaso et al., 2010) in Python 2.7. The total sequences were clustered using UCLUST into OTUs tables (operational taxonomic units) using the Greengene database (GG 13_8_otus) as a reference with a range of 97% of similarity and using the closed system with the command pick_closed_reference_otus.py. Taxonomy summaries including relative abundance data were generated using summarize_taxa.py, plot_taxa_summary.py and plot_taxa_through_plots.py commands. In all the cases, we used the data filtering option of 0.01% in abundances because it is reported that filtering database decreases the estimation error (Kuczynski et al., 2012; Navas-Molina et al., 2015).

Alpha diversity was evaluated using the function of alpha_rarefaction.py from QIIME, that calculate alpha diversity on each sample in an OTUs table, using a variety of alpha diversity metrics as Shannon-Wiener index, Simpson index, Otus_observed and Chao1 value. Each metrics result was analyzed by ANOVA applying the Tukey-Kramer test (0.95 confidence interval) to estimate the significance difference between the samples. Beta diversity was calculated by beta_diversity_through_plots.py, a workflow script for computing beta diversity distance matrices (UniFrac unweighted method) and generating Principal coordinates analysis (PCoA) plots from QIIME.

The normalized OTUs table (0.01% abundance filter) was used to estimate functional features present in the samples, using PICRUSt (Phylogenetic Investigation of Communities by Reconstruction of Unobserved States) version 1.1.0 (Langille et al., 2013) and the Greengenes databases 16S_13_5 and KO_13_5. The OTUs table was normalized to obtain the metagenomic functional predictions at different hierarchical KEGG levels using normalize_metagenomes.py, predict_metagenomes.py and categorize_by_function.py scripts of the same software.

For the statistical studies of the functions, we used STAMP (Statistical analysis of taxonomic and functional profiles) version 2.1.3, through ANOVA analysis applying the Tukey-Kramer test (0.95 confidence interval) to evaluated gene abundance of each function. R statistical program (version 3.3.3) was used to make plots using “ggplot2” and “dplyr” libraries.

## Results

### Sample composition

To determine the microenvironmental conditions that affect the microbial communities and metabolic characteristics of the fermentation process of Tuba, we measured the sugars (sucrose, glucose, and fructose), water-soluble proteins, ethanol and acetate concentrations, as well as the pH value, at different fermentation times (Supplementary Table 1S). The starting sap from palm 1 (Tuba P1) was the sample with the highest concentration in glucose and fructose with 61.4 and 47.3 g/L respectively at 12 hours, 4.7% (v/v) in ethanol and 6.0 g/L in acetate at 35 hours (**Figure 2A**). Tuba P2 was the sample with the lowest concentration of monosaccharides at the beginning of the fermentation and high sucrose concentration (121.7 g/L), however, at the last fermentation time, the acetate and ethanol concentrations were 3.5 g/L and 0.6% (v/v) respectively (**Figure 2B**). The glucose and fructose concentration in Tuba P3 were 39.8 and 29.1 g/L respectively at 12 hours. Having at the end of the fermentation the highest concentration of ethanol (5% v/v) in comparison with all samples (**Figure 2C**). The pH values of the Tuba P1, P2 and P3 decreased from 3.7 to 2.8 during the fermentation process. The water-soluble protein concentration of the Tuba samples showed low values from 0.06 to 0.1 g/L. In the case of the commercial Tuba samples, all samples presented similar composition values, (average values: 40.5 g/L of sucrose, 40.0 g/L of glucose, 42.53 g/L of fructose, 1.6 g/L of acetate, 0.1% (v/v) of ethanol and pH of 4 **Figure 2D**).

**Table 1.** Alpha diversity values.

| Sample | Alpha diversity metrics |  |  |  |
| --- | --- | --- | --- | --- |
|  | Chao1 index | Observed_otus | Shannon index | Simpson index |
| **Comercial Tuba**<br>**(n=6)** | 49.240 ± 3.758 | 42.600 ± 5.609 | 2.619 ± 0.006** | 0.692 ± 0.002** |
| **Tuba P1 (n=6)** | 49.715 ± 5.307 | 42.508 ± 6.621 | 2.876 ± 0.010** | 0.781 ± 0.001** |
| **Tuba P2 (n=6)** | 35.863 ±<br>4.654** | 26.588 ± 5.235* | 1.595 ± 0.011** | 0.491 ± 0.002** |
| **Tuba P3 (n=6)** | 49.559 ± 5.305 | 41.229 ± 7.006 | 2.468 ± 0.014** | 0.704 ± 0.001** |

\* show significant difference with $p<0.01$ . \*\* show significant difference with $p<0.001$

**Figure 2.**
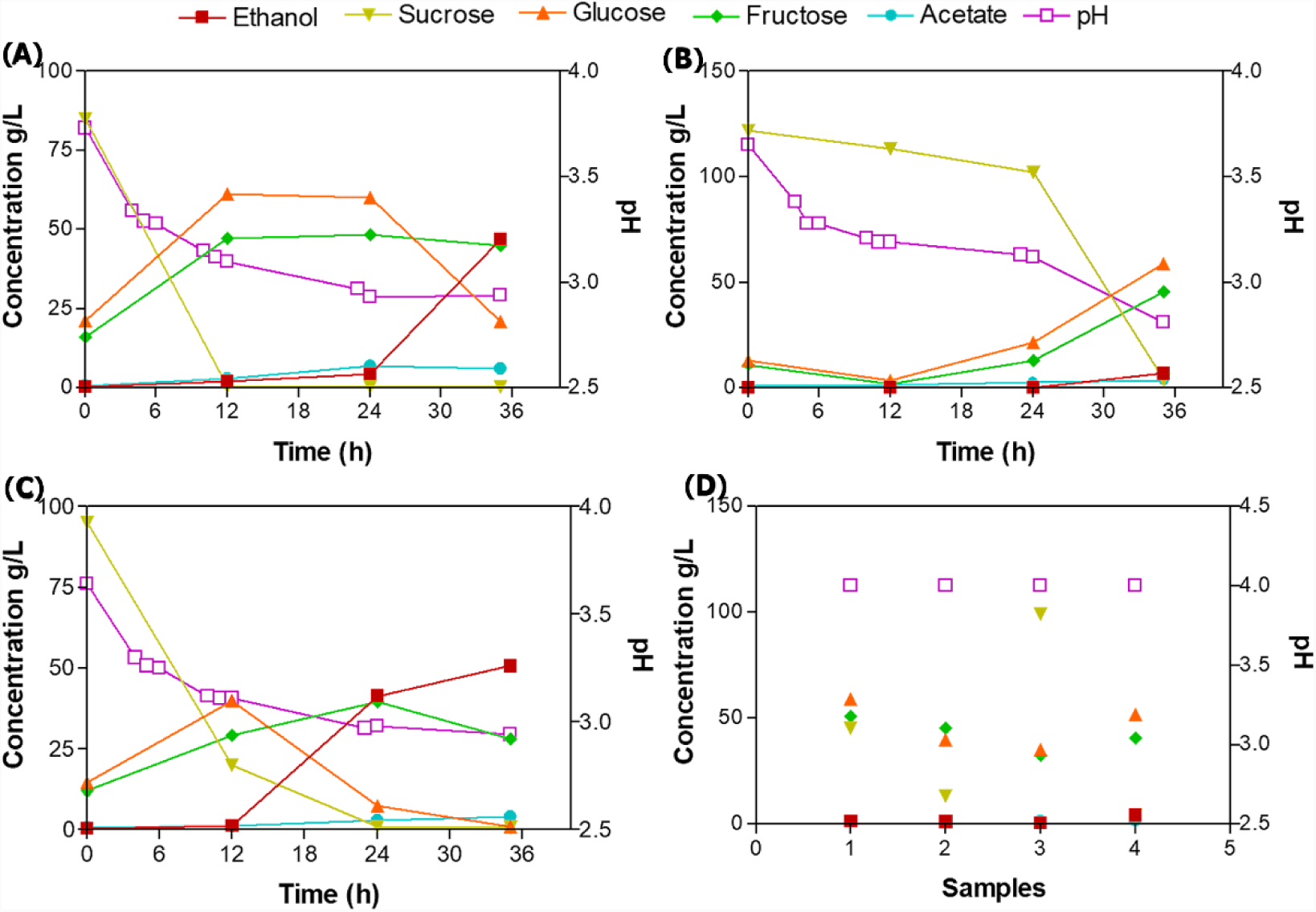
Composition of the laboratory fermented Tuba and commercial Tuba. A) Tuba P1, B) Tuba P2, C) Tuba P3 and D) Commercial Tuba samples. Each number correspond to one sample. Right axis represented pH value.

### Taxonomic classification

A total of 302,398 sequences were obtained from the 16S amplicon libraries of 3 independent fermentation experiments samples at four different times, and four different commercial samples from different local vendors. Different fermentation times and commercial samples, with an average of 75,594 ±587 sequences per Tuba samples. A total of 123 OTUs were detected in all Tuba samples. However, filtering database with 0.01% relative abundance filter, the OTUs were reduced to 28 as the more abundant. The taxonomic identification was elaborated using the last filter mentioned and the top 10 most abundant genera of the 16 samples are showed in **Figure 3**. The genera that predominate in all the samples were *Fructobacillus, Leuconostoc, Gluconacetobacter*, *Sphingomonas*, *Vibrio* and some genera of the Enterobacteriaceae family. Additionally, analyzing the Enterobacteriaceae populations with the lower abundance we found genera as *Erwinia, Klebsiella, Serratia,* and *Cronobacter (*Supplementary Figure 1S). The population dynamic had a similar trend in Tuba fermented in controlled conditions but with the different percentage in the OTUs abundances; we observed a decrease of *Vibrio* genus and an increase of lactic acid bacteria (LABs), acetic acid bacteria (AABs) and some proteobacteria as *Sphingomonas* through the fermentation time (**Figure 3**).

**Figure 3.**
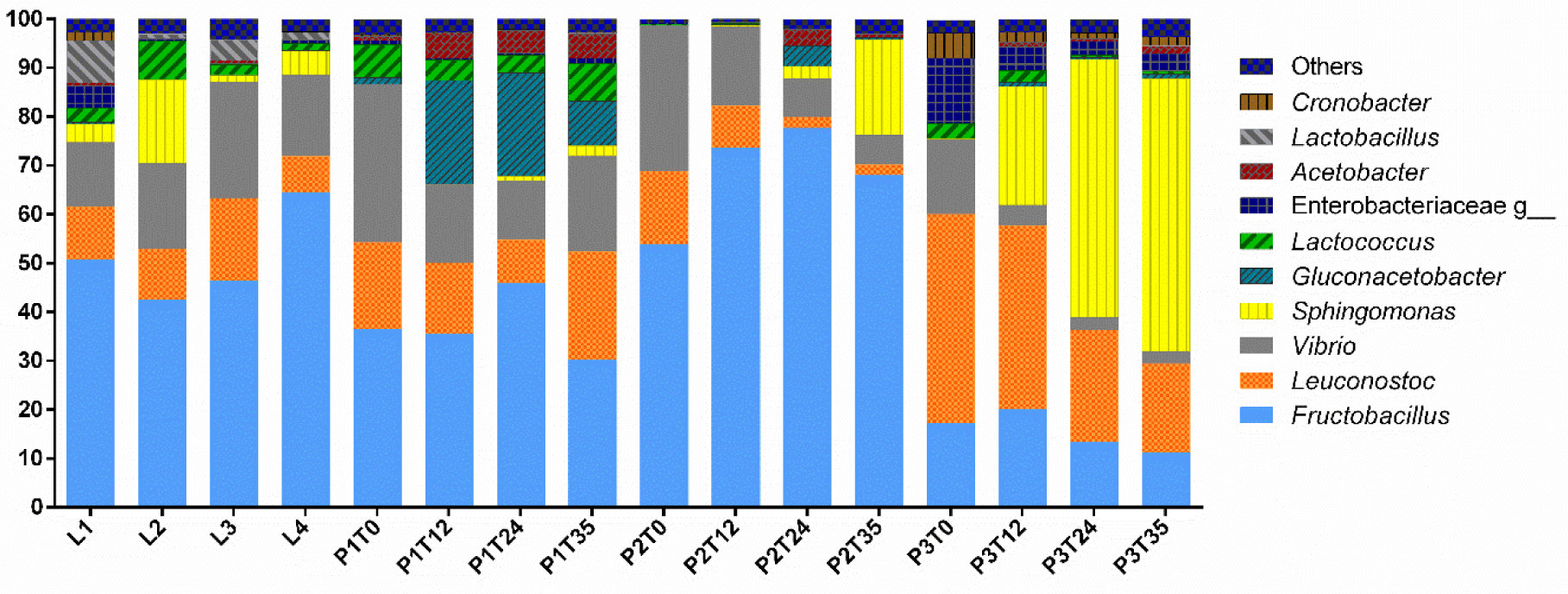
Taxonomic identification. The graph represented the top ten genera using the 0.01% abundance filter OTUs table.

### Diversity analysis

The analysis as richness (observed_otus), dominance (Simpson), equity (Shannon index) and singletons (Chao1 value), which reflect the Alpha diversity in bacteria communities, showed significant differences between all Tuba samples (**Table 1**) (Supplementary Figure 2S). Tuba P1 was the most diverse with the highest values in the four diversity index, then Tuba P3 and commercial Tuba samples had similar index values, and finally, Tuba P2 was the least diverse with the lowest values. After of ANOVA statistical analysis, we found that in Chao1 and Observed_otus tests Tuba P2 was the only showing significant difference. Nevertheless, in Shannon and Simpson index the four groups showed significant difference among each other (**Table 1**).

The Unweighted UniFrac distance using the 0.01% abundance filter, did not show clustering by fermentation time (**Figure 4A**). However, a clustering was observed by sample origin (**Figure 4B**). In the graphic of origin of the sample we also observe a grouping by quadrant of the all the Tuba samples, however, Tuba P2 showed the greatest dispersion in the data, which indicated a big difference between the fermentation times in Tuba P2. A similar effect is observed in Tuba P1 where two fermentation times (0 h and 35 h) show similar beta diversity values compared to commercial samples and Tuba P3. Otherwise, the samples, which were in the same quadrant as Tuba P3 and the commercial Tuba, were considered strongly related (**Figure 4**).

**Figure 4.**
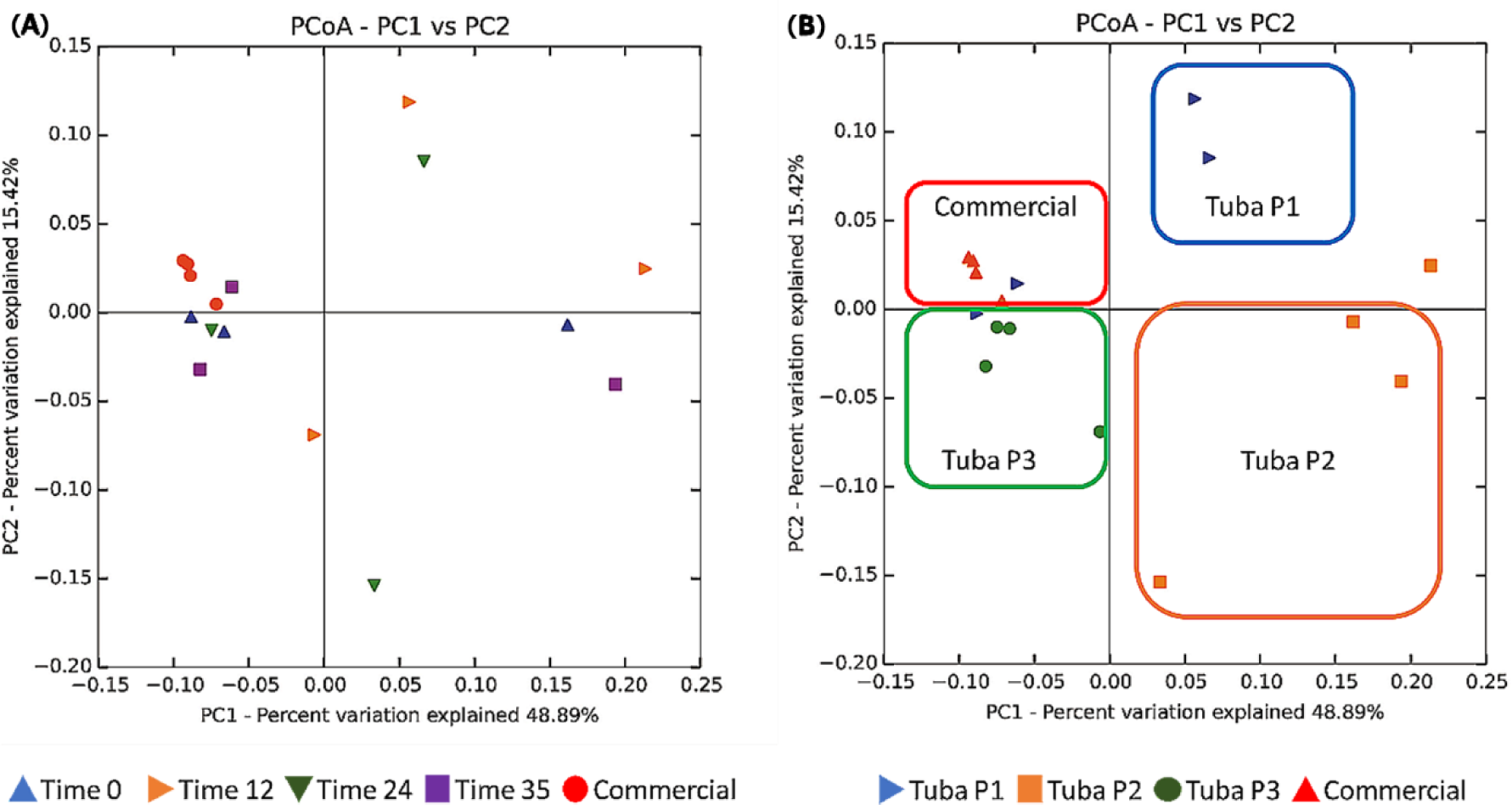
Beta diversity. A) Associate with respect to the fermentation time. B) Associate with respect to the origin of the sample. Analysis performed by the Unifrac unweighted technique with 0.01% abundance filter and plotted with the Principal Coordinates Analysis (PCoA). The color boxes show a manual grouping of the data.

### Functional profile prediction

To understand the functionality of the bacterial community in Tuba fermentation, PICRUSt software was used to predict the metagenomic profiles of the samples. Initially, we obtained functional characteristics of the 3 KEGG levels (Level 1: general cellular functions, Level 2: Specific functions i.e. carbohydrates metabolism, and Level 3: Specific pathway associated with a specific function) (http://www.genome.ad.jp/kegg/). We limited our analysis to the level 3 and we discarded elementary cellular functions such as replication, translation, and functions associated with human diseases (cancer) or poorly characterized functions, to analyze specific genes related with functions of biotechnological relevance. Considering the 328 registered functions on KEGG, we found the 19 most abundant functions were associated with carbohydrates metabolic process, vitamins, amino acid, antibiotics and antioxidant molecules biosynthesis (**Figure 5A**), suggesting that the production of those compounds may be taking place during Tuba fermentation. Many of those functions are common to fermented foods, however some of them are not obvious and by this analysis we can attribute the function to a specific OTU. As an example *Sphingomonas* and *Gluconacetobacter* had more abundance percentage in the enzyme *15-cis-phytoene synthase (***Supplementary Figure 4S**). After an ANOVA test, we found functions without a significant difference as the carotenoid biosynthesis (**Figure 5B**), this means that no matter what is the sample origin, this function may have present at the same gene abundance in the four groups. Otherwise, there were functions with a significant difference, such as peptidases biosynthesis that had more gene abundance in Tuba P2 samples (**Figure 5C**). Each sample had more abundance in genes associated with a specific function, for example, antioxidant, antibiotic compounds, and folate biosynthesis in Tuba P3, lipopolysaccharide and lysine biosynthesis genes in Tuba P1, finally the 4 commercial Tuba samples may have bacteria with genes associated mainly with folate biosynthesis and peptidases.

**Figure 5.**
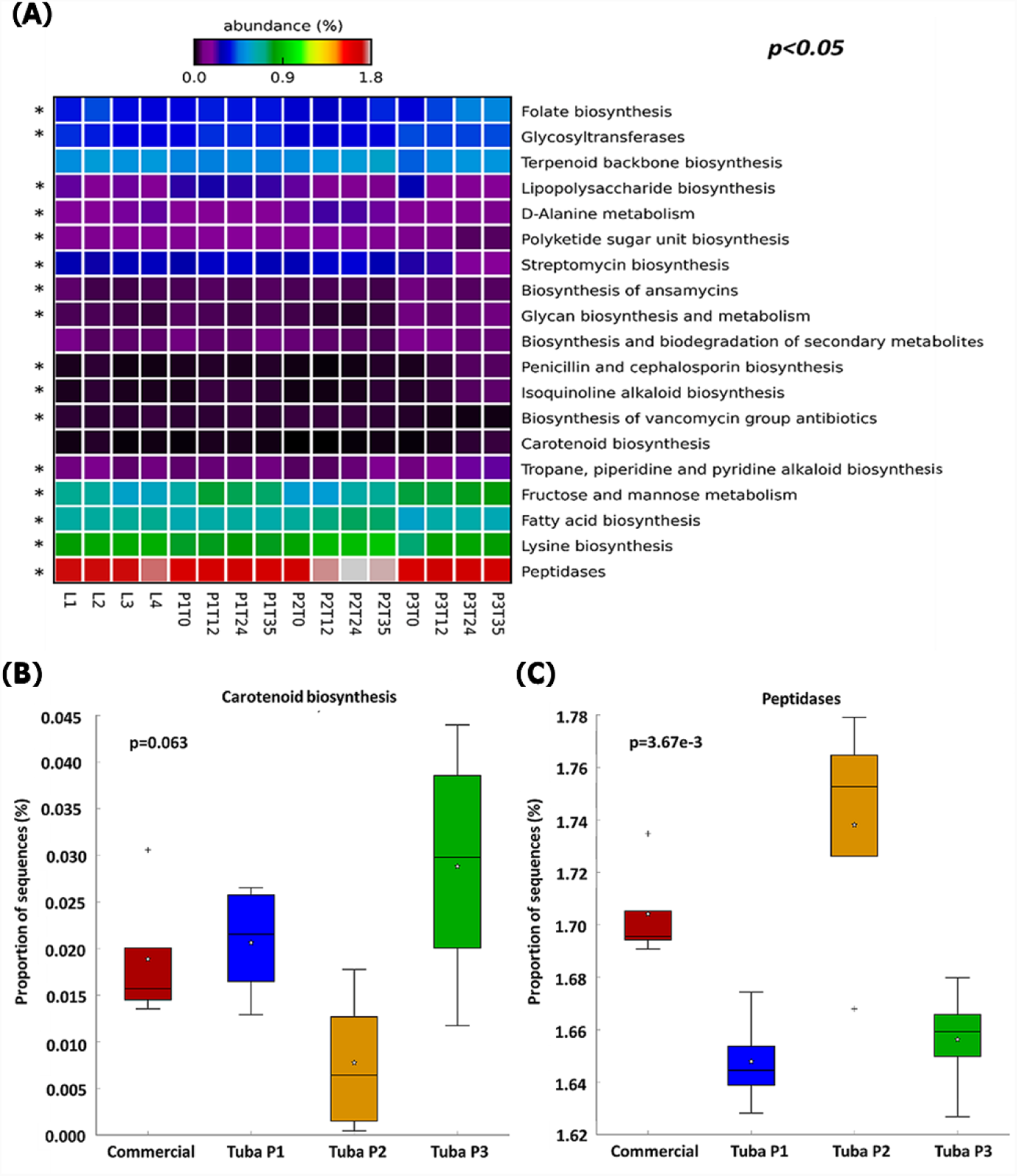
Abundance of sequences associate with functions. An ANOVA was performed with Tukey-Kramer (0.95). A) Heatmap of the percentage of genes associated with functions, discarding elementary cellular functions. Asterisk show functions with considerable difference (*p<0.05*). B) Box plot of not significant difference function. C) Box plot of a function with significant difference.

## Discussion

In this study, we were interested in associating abiotic variables such as the initial sap composition to the bacterial diversity, population dynamics, and functions of Tuba. All sap samples had a high initial sugar content, however, they showed significant variations (Supplementary Table 1S). Fermentation profile was similar among samples Tuba P1 and Tuba P3 and different in Tuba P2 (**Figure 2**). Tuba P1 and P3 started to ferment immediately, at 12 h we found acetate and ethanol production; and sugar hydrolysis to glucose and fructose. Samples from Tuba P2 showed a different behavior in which the sucrose hydrolysis and ethanol production were delayed. We may attribute this behavior to two possible (not mutually exclusive) explanations. A) The inhibitory nature of sucrose on its own hydrolysis (Goosen et al., 2007) since Tuba P2 was higher initial sucrose load, B) a lower abundance of yeasts. Here we attribute all the ethanol production during the fermentation to yeasts since we did not find any ethanologenic bacteria, as it has been found in other ethanolic fermented beverages (Díaz-Ruíz et al., 2003; Escalante et al., 2008). From the analyzed bacterial population we may attribute the sucrose hydrolysis to *Fructobacillus* and *Leuconostoc* genera, through the functional profiling we found that those genera carry the beta-fructofuranosidase gene responsible for sucrose hydrolysis (Supplementary Figure 3S).

The aim of this study was to find the bacterial composition of Tuba and its dynamics and function during the production of the fermented beverage. We found that main bacterial genus present in all samples were LABs (*Fructobacillus*, *Leuconostoc,* and *Lactococcus*), AABs (*Gluconacetobacter* and *Acetobacter*) and Proteobacteria (*Vibrio*). Regarding the diversity indexes calculated here, we showed that all Tuba samples (laboratory produced and commercial) had the same 10 main genera but in different abundance (**Figure 3**), having significant differences in Shannon and Simpsons indexes (**Table 1**). Using the Observed OTUs and Chao1 indexes, the only sample with a significant difference is Tuba P2, its significant lower value (p-value <0.001), this means a lower number of bacterial genus and high dominance in the sample. Beta diversity indicated that the origin of the sample as the best parameter for grouping, not the fermentation time (**Figure 4**). This showed that biotic and abiotic factors conditions (seasonality, plant physiology, age, soil conditions, and other abiotic variables such as water irrigation and other environmental factors) affected the bacterial diversity (Coleman-Derr et al., 2016; Fonseca-García et al., 2016; Staley et al., 2014) at different times of the fermentation. It is worth mentioning that the sap samples were collected after they were harvested by the producer and we took all the precautions to conserve the initial bacterial community.

Palm wine is a worldwide consumed beverage with different characteristics mainly depending on palm species used and its production process. Our findings found that southern pacific Mexican commercial Tuba has low alcohol content; even as we showed, it can reach higher concentration if fermented for a longer time (more than 35 h at specific conditions, i.e. P3T35). In Guerrero, Tuba is consumed as a refreshing, hydration drink and it is empirically used as a traditional aid for gastrointestinal discomfort and nutritional aid (De la Fuente-Salcido et al., 2015; Velázquez-Monreal et al., 2011). The functional profile prediction (**Figure 5**) enable us to support to the empirical uses of Tuba. Some functions such as antibiotic production pathways and peptidases genes have been related to its uses for stomach discomfort, which support the anti-pathogenic and probiotic use this fermented beverages as it has been studied elsewhere for other plant sap derived fermented beverages (Escalante et al., 2016). We also found a prediction of bacterial production of antioxidant compounds such as terpene and carotenoid biosynthesis pathways. Plant products are known to be rich in antioxidants in specific coconut water (*Cocos nucifera* L. variety Chandrasankara) has been tested for its ability to scavenge free radicals, and they found a good antioxidant activity percentage (Mantena et al., 2003), the bacterial production of antioxidant compounds (as the enzyme *15-cis-phytoene synthase* that is related with the phytoene production) predicted here adds to the already know antioxidant activity. In terms of nutritional properties, we found the folate and amino acid biosynthesis pathways, which together with the known properties of coconut sap/water (Debmandal and Mandal, 2011; Manivannan et al., 2016) makes Tuba functional fermented beverage. Significant differences in the predicted functions can be related to the OTUs abundance; as a perspective we may study the metatranscriptome of our samples to find the gene expression in the microbial community. Therefore, our findings may guide future efforts to create palm fermented beverages with specific functional properties.

In this work, we reported for the first time the bacterial diversity and potential functional analysis through the fermentation process of the Tuba. With the knowledge of microbiota diversity and metabolic functional inference, the Tuba production can be controlled by adjusting the presence and abundance of beneficial genera that contributes with the functional characteristics of the Tuba. It also contributed to establishing the microbiological basis of its empirical uses. Additionally, the isolation of bacteria from these samples may provide us with new species with probiotic potential.

## Acknowledgments

We would like to acknowledge Raunel Tinoco Valencia for HPLC analysis. Filiberto Sánchez Lopez for technical assistance in library preparation. Funding from DGAPA-PAPIIT-UNAM IA201518. Fernando Astudillo-Melgar held a CONACyT scholarship with register number 597135.

## Supplementary material Figures

**Figure 1S.**
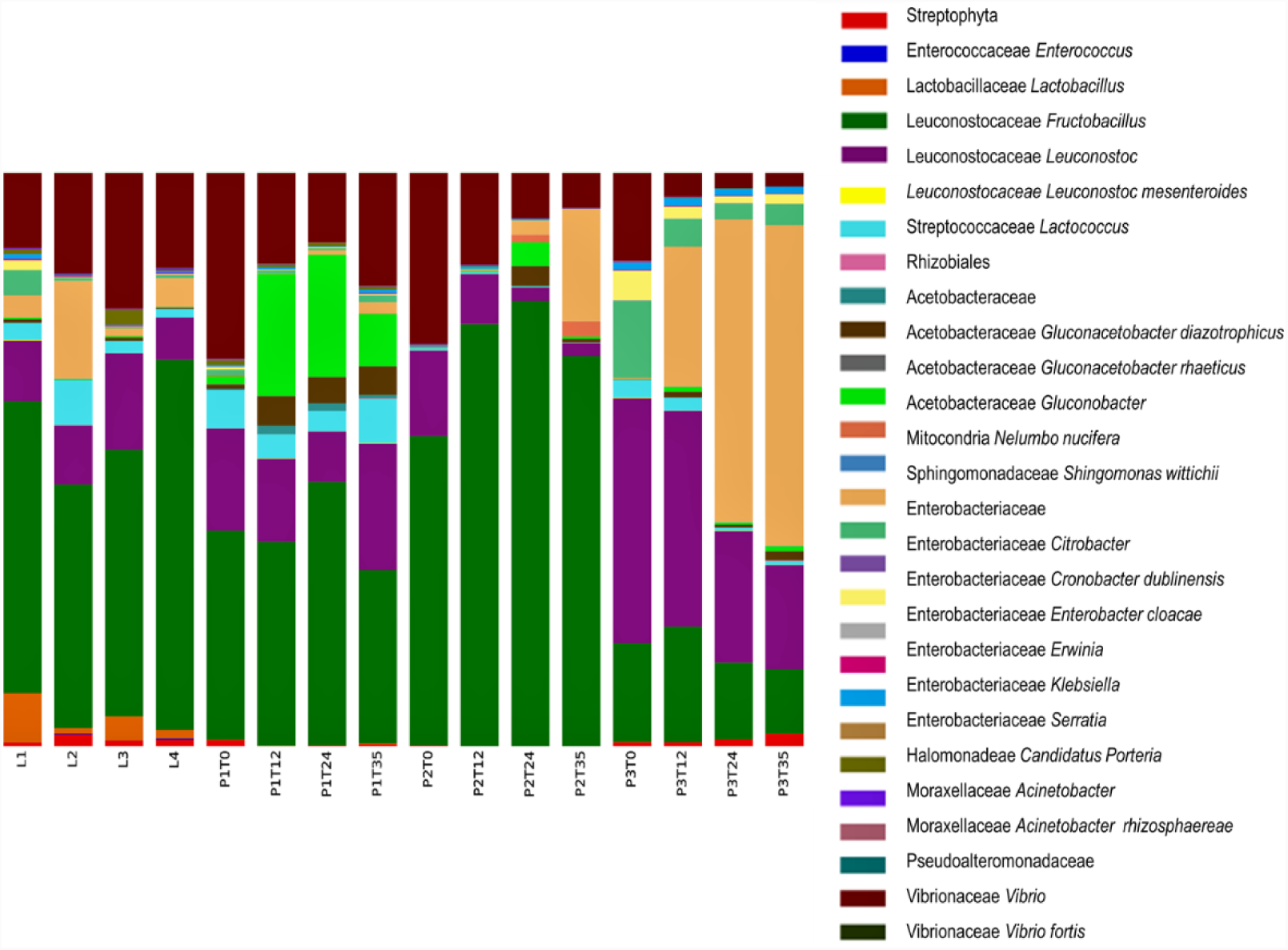
Taxonomic identification. Most abundant OTU’s using the 0.01% abundance filter OTUs table.

**Figure 2S.**
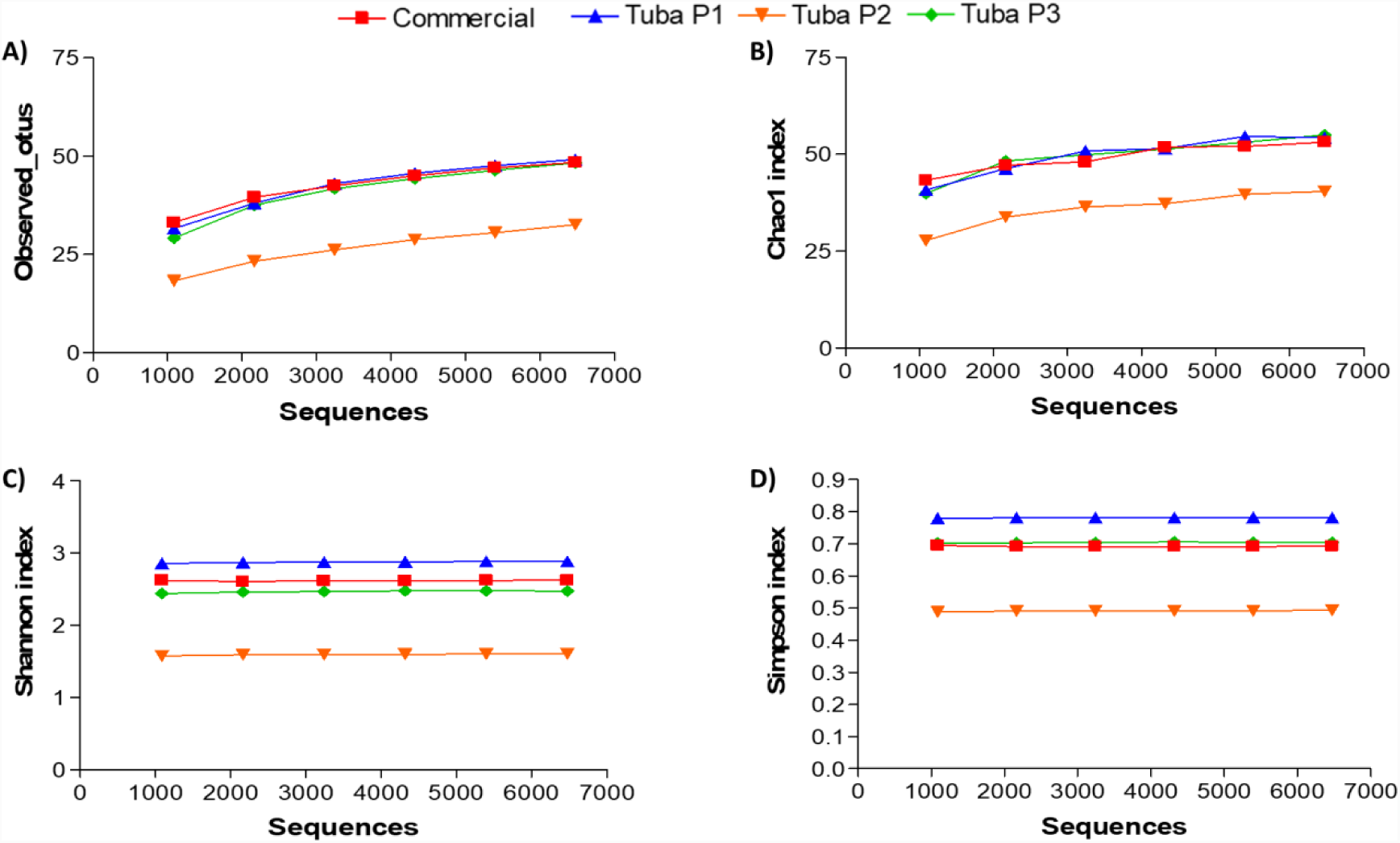
Alpha diversity rarefaction plots with 0.01%. A) Observed_otus, B) Chao1, C) Shannon and D) Simpson. Each population is represented for a specific color in all the graphics.

**Figure 3S.**
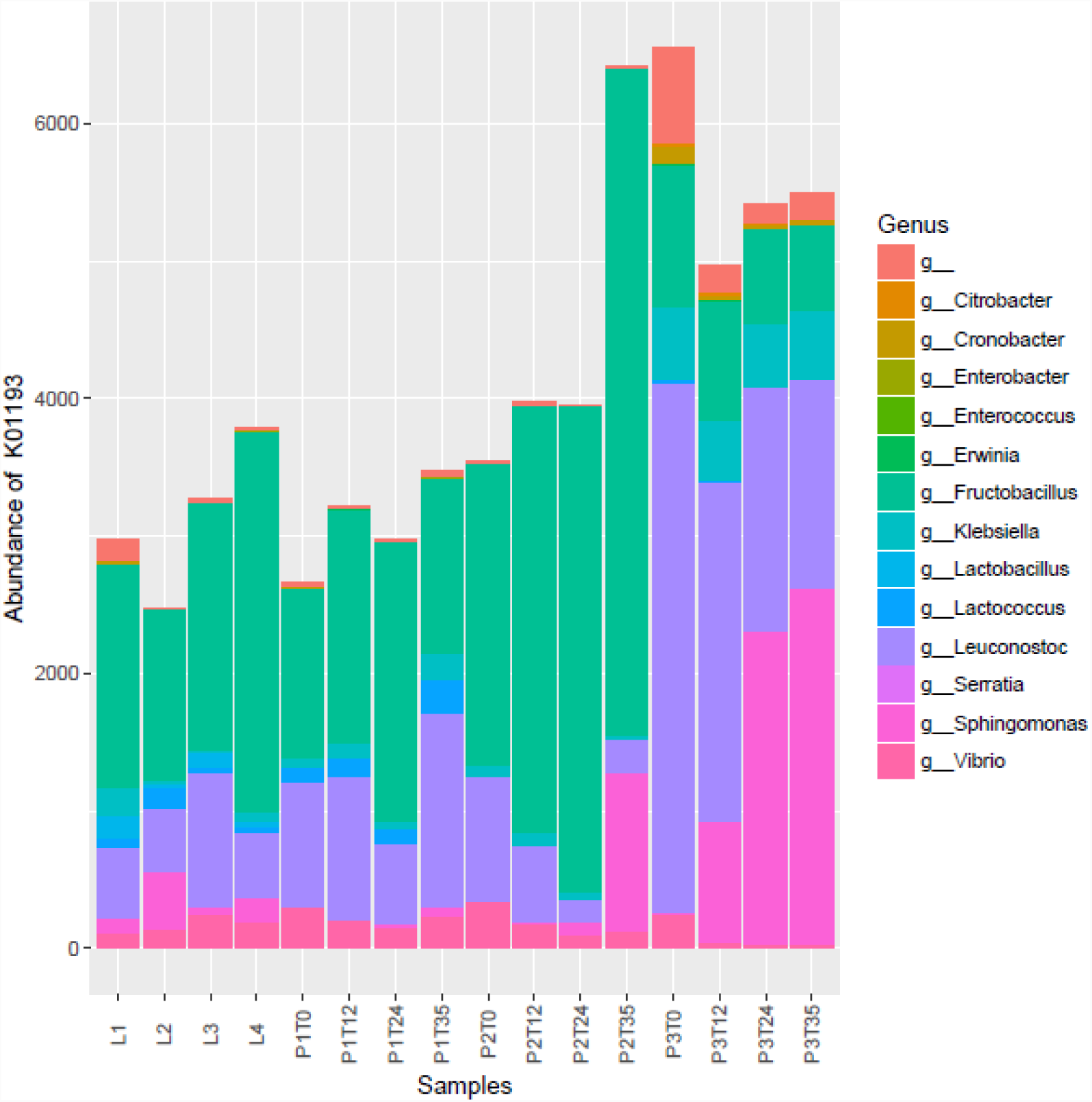
Abundance of invertase gene (K01193). Analysis performed with the function “metagenome_contributions.py” obtained by PICRUSt analysis and plotted with R studio.

**Figure 4S.**
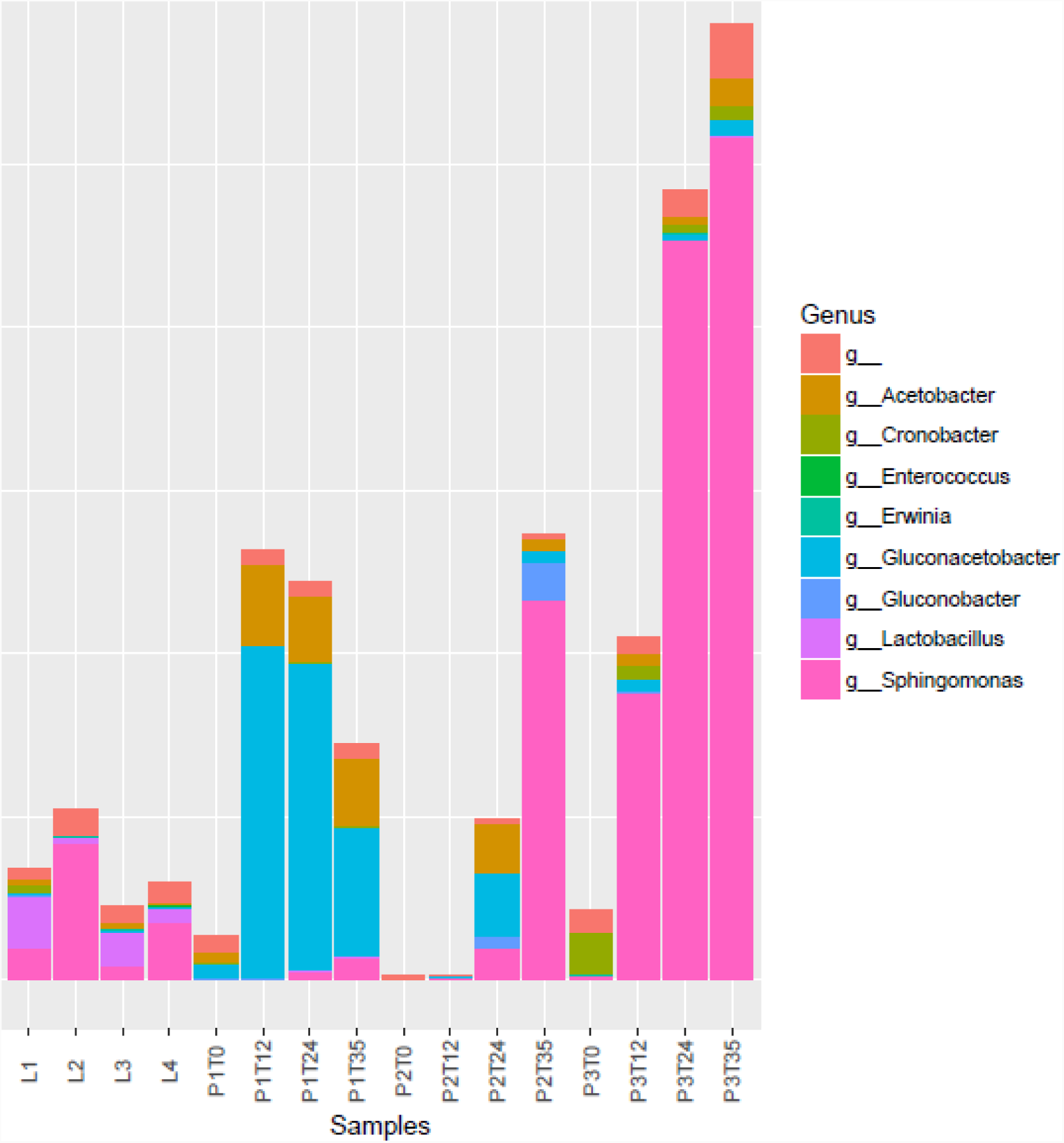
Main bacteria with 15-cis-phytoene synthase gene (K02291 KEGG code). Analysis performed with the function “metagenome_contributions.py” obtained by PICRUSt analysis and plotted with R studio.

**Table 1S.**
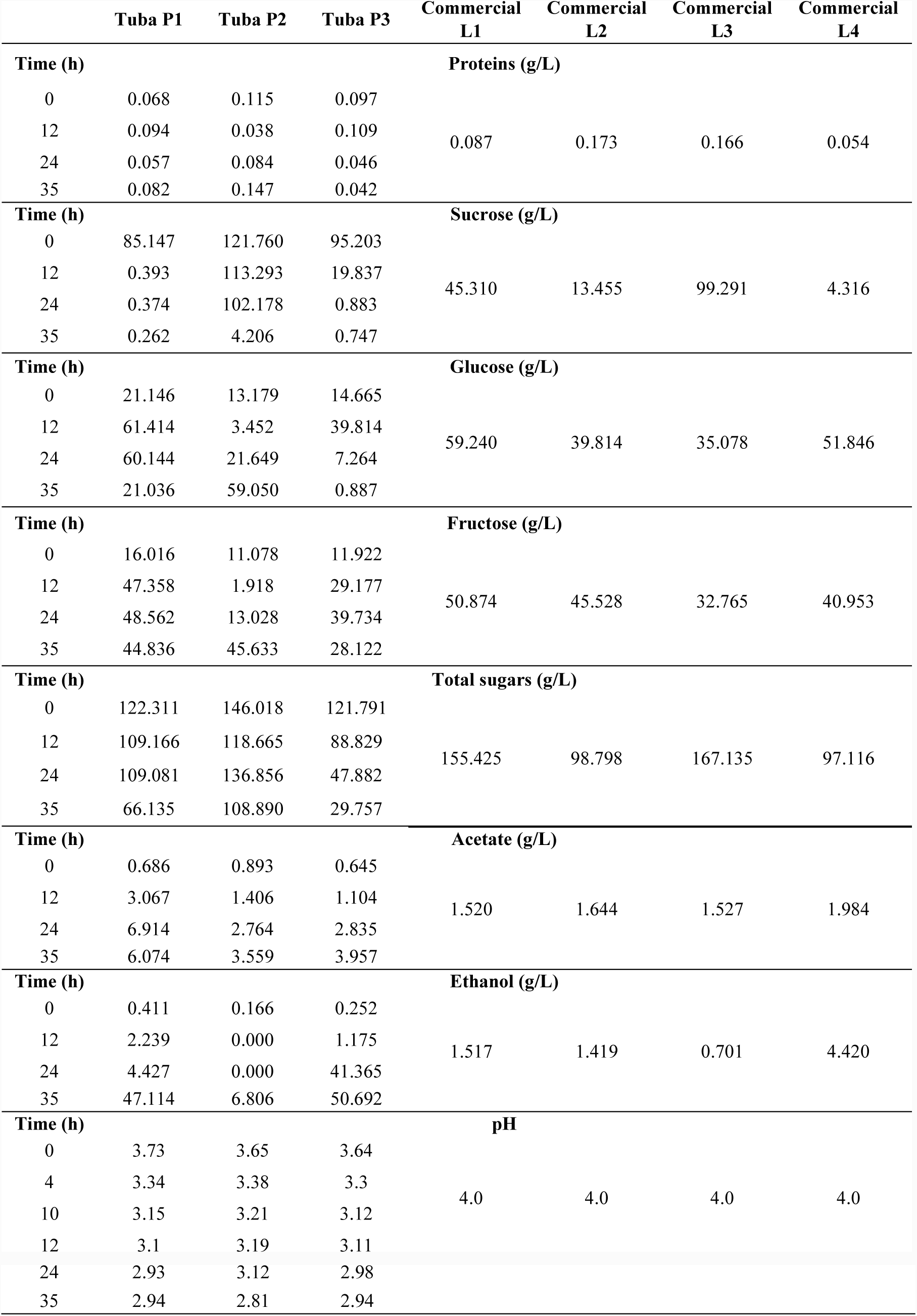
Chemical composition of the Tuba.

|  | **Tuba P1** | **Tuba P2** | **Tuba P3** | **Commercial**<br>**L1** | **Commercial**<br>**L2** | **Commercial**<br>**L3** | **Commercial**<br>**L4** |
| --- | --- | --- | --- | --- | --- | --- | --- |
| **Time (h)** | **Proteins (g/L)** |  |  |  |  |  |  |
| 0 | 0.068 | 0.115 | 0.097 |  |  |  |  |
| 12 | 0.094 | 0.038 | 0.109 | 0.087 | 0.173 | 0.166 | 0.054 |
| 24 | 0.057 | 0.084 | 0.046 |  |  |  |  |
| 35 | 0.082 | 0.147 | 0.042 |  |  |  |  |
| **Time (h)** | **Sucrose (g/L)** |  |  |  |  |  |  |
| 0 | 85.147 | 121.760 | 95.203 |  |  |  |  |
| 12 | 0.393 | 113.293 | 19.837 | 45.310 | 13.455 | 99.291 | 4.316 |
| 24 | 0.374 | 102.178 | 0.883 |  |  |  |  |
| 35 | 0.262 | 4.206 | 0.747 |  |  |  |  |
| **Time (h)** | **Glucose (g/L)** |  |  |  |  |  |  |
| 0 | 21.146 | 13.179 | 14.665 |  |  |  |  |
| 12 | 61.414 | 3.452 | 39.814 | 59.240 | 39.814 | 35.078 | 51.846 |
| 24 | 60.144 | 21.649 | 7.264 |  |  |  |  |
| 35 | 21.036 | 59.050 | 0.887 |  |  |  |  |
| **Time (h)** | **Fructose (g/L)** |  |  |  |  |  |  |
| 0 | 16.016 | 11.078 | 11.922 |  |  |  |  |
| 12 | 47.358 | 1.918 | 29.177 | 50.874 | 45.528 | 32.765 | 40.953 |
| 24 | 48.562 | 13.028 | 39.734 |  |  |  |  |
| 35 | 44.836 | 45.633 | 28.122 |  |  |  |  |
| **Time (h)** | **Total sugars (g/L)** |  |  |  |  |  |  |
| 0 | 122.311 | 146.018 | 121.791 |  |  |  |  |
| 12 | 109.166 | 118.665 | 88.829 | 155.425 | 98.798 | 167.135 | 97.116 |
| 24 | 109.081 | 136.856 | 47.882 |  |  |  |  |
| 35 | 66.135 | 108.890 | 29.757 |  |  |  |  |
| **Time (h)** | **Acetate (g/L)** |  |  |  |  |  |  |
| 0 | 0.686 | 0.893 | 0.645 |  |  |  |  |
| 12 | 3.067 | 1.406 | 1.104 | 1.520 | 1.644 | 1.527 | 1.984 |
| 24 | 6.914 | 2.764 | 2.835 |  |  |  |  |
| 35 | 6.074 | 3.559 | 3.957 |  |  |  |  |
| **Time (h)** | **Ethanol (g/L)** |  |  |  |  |  |  |
| 0 | 0.411 | 0.166 | 0.252 |  |  |  |  |
| 12 | 2.239 | 0.000 | 1.175 | 1.517 | 1.419 | 0.701 | 4.420 |
| 24 | 4.427 | 0.000 | 41.365 |  |  |  |  |
| 35 | 47.114 | 6.806 | 50.692 |  |  |  |  |
| **Time (h)** | **pH** |  |  |  |  |  |  |
| 0 | 3.73 | 3.65 | 3.64 |  |  |  |  |
| 4 | 3.34 | 3.38 | 3.3 | 4.0 | 4.0 | 4.0 | 4.0 |
| 10 | 3.15 | 3.21 | 3.12 |  |  |  |  |
| 12 | 3.1 | 3.19 | 3.11 |  |  |  |  |

|  |  |  |  |
| --- | --- | --- | --- |
| 24 | 2.93 | 3.12 | 2.98 |
| 35 | 2.94 | 2.81 | 2.94 |

